# An atlas-scale generative model for unified representation learning of bulk RNA-seq data

**DOI:** 10.64898/2026.06.18.733198

**Authors:** Amit Pande, Bora Uyar, Altuna Akalin

## Abstract

Public bulk RNA-seq repositories contain hundreds of thousands of samples, creating opportunities for large-scale representation learning, but integration across studies remains challenging because of heterogeneous annotations, experimental protocols, and technical variation. While pre-trained foundation models are now widely available for single-cell RNA-seq, comparable resources for bulk RNA-seq remain scarce, motivating a model that learns a unified, tissue-aware representation directly from bulk data. We trained a supervised variational autoencoder (VAE) on a compendium of 118,263 bulk RNA-seq samples that we assembled from TCGA, GTEx, and ARCHS4 and mapped to 42 tissue categories. The model classifies tissue of origin at 94.9% balanced accuracy (weighted F1 96.2%) and compresses 16,115 genes into a 121-dimensional latent space. Tissue identity is the primary organizing axis of the latent space, while source effects remain secondary. To assess the impact of data volume, we constructed training sets at three different scales (38K, 75K, and 118K samples). Our results demonstrated that reconstruction fidelity improved incrementally with each expansion of the dataset, but with diminishing returns. We validated the model on an independent cohort of 734 paediatric tumour samples from TARGET, achieving 84.6% agreement with the expected tissue of origin. The trained model and code are available at GitHub (https://github.com/BIMSBbioinfo/flexynesis_tissue_vae_manuscript) with an interactive web application.

## Introduction

Tissue identity is a dominant source of biological variation in bulk RNA-seq, and large public archives — GEO, TCGA, GTEx, and the cancer cell-line encyclopedias — collectively contribute expression profiles for hundreds of thousands of human samples (1–5). The bottleneck is no longer data generation but interpretation: samples come from different laboratories, on different platforms, processed through different pipelines, and annotated with different vocabularies.

Conventional approaches to this heterogeneity — ComBat, limma’s removeBatchEffect, Harmony — treat source variation as a confounder to be removed (6–8). These methods work well when source labels are clean and the underlying biology is consistent across studies. In experimental contexts that differ in genuinely biological ways, batch correction can remove real signals alongside technical noise.

Single-cell foundation models such as scGPT, Geneformer, and scBERT capture biological structure from large transcriptomic datasets (9–11), but they are built for single-cell data and address a different problem. In bulk RNA-seq, each sample is a composite of cell types, and the variation of interest sits at the tissue level rather than the cellular level.

Compared with the rapidly expanding ecosystem of single-cell foundation models, bulk RNA-seq has comparatively few large-scale representation-learning resources of this kind: a representation that captures tissue identity, accommodates source heterogeneity, and does not require foundation-model-scale resources. The most prominent recent effort is BulkFormer, a transcriptome-wide foundation model pretrained on over 500,000 bulk profiles, which we use as the benchmark for our comparisons (12). For this kind of model, data quality matters more than quantity. A recent benchmark varied pretraining data size and diversity for single-cell foundation models and found that accuracy plateaued below current dataset sizes, with no evidence of the data scaling observed in language models; on several tasks, simple baselines did just as well (13). Tissue of origin is a well-defined target, and for such targets the supervision signal matters more than raw data volume. We therefore use a variational autoencoder with a supervised head: the encoder compresses expression into a small latent space, and the tissue label shapes that space by tissue identity. A representation of this kind has direct practical utility: it provides a common coordinate system for comparing samples across studies, enables tissue-of-origin quality control for large or mislabelled archives, and yields a compact feature space that downstream models can reuse without per-study batch correction.

In this work, we train a supervised variational autoencoder to learn such a representation. The model is trained on 118,263 bulk RNA-seq samples drawn from three public repositories, mapped to 42 tissue categories using the UBERON ontology (14). We use this representation to ask whether it separates tissues, withstands missing genes, and transfers to an unseen paediatric cancer cohort (TARGET). We observe that the latent space is organized chiefly by tissue identity rather than data source, reconstructs expression robustly, and maps held-out tumours to their developmental tissue of origin, giving a reusable, tissue-aware representation of bulk RNA-seq.

## Results

### Building a tissue-curated, balanced multi-source bulk RNA-seq compendium

The first task was to assemble a training set that is large, consistently annotated, and biologically representative of primary tissues. ARCHS4 is the largest single source and also the most heterogeneous: it re-processes raw GEO submissions uniformly, so its 888,821-sample human archive contains a mixture of bulk and single-cell RNA-seq, cancer cell-line profiles, and a wide range of experimental platforms (5). We used the ARCHS4 single-cell probability score to keep bulk-only samples (score < 0.5), reducing the pool to 605,614 samples.

From this pool, 411,318 samples were extracted and their free-text tissue annotations were mapped to the UBERON ontology by keyword matching followed by manual curation, yielding 212,412 tissue-labelled samples. These were combined with TCGA (n = 9,400 tumour samples) and GTEx (n = 14,768 normal tissues), forming a candidate pool of 236,580 samples; 10 samples with unmappable tissue annotations were removed, leaving 236,570. We then applied two filters: ARCHS4 cell-line-flagged samples were excluded by ENCODE/CCLE-style identifier matching, reducing the pool to 188,790 and each UBERON category was capped at 10,000 samples, yielding the final compendium of 146,537 samples (118,263 training / 28,274 test, stratified by tissue category) across 42 tissue categories. The final compendium comprises 123,127 ARCHS4, 14,072 GTEx, and 9,338 TCGA samples; DepMap is absent by design (see Limitations) (Table 1).

**Table 1.** Multi-source filtering and curation funnel from raw public archives to the final 118,263-sample training compendium. ARCHS4 contributes the bulk of input heterogeneity and requires the largest reduction (888K → 118K, ∼13× compression) to reach the final curated state. TCGA, GTEx, and DepMap enter the pool at their full carried-over sizes; the cell-line filter (step 9) removes all DepMap samples (1,451) together with ARCHS4 cell-line-flagged samples, so DepMap is absent from the final compendium by design (see Limitations). The final compendium comprises 123,127 ARCHS4, 14,072 GTEx, and 9,338 TCGA samples (146,537 total); per-source counts fall below their pool-entry values where the 10,000-per-tissue cap applied.

| Step | Source / Operation | Output samples | Cumulative effect |
| --- | --- | --- | --- |
| 1 | ARCHS4 v2.5 (all human RNA-seq submissions) | **888,821** | Raw GEO pool — bulk + single-cell + cell-line + mixed platforms |
| 2 | ARCHS4 → bulk-only filter (single-cell prob. < 0.5) | **605,614** | Single-cell submissions removed (~32% reduction) |
| 3 | ARCHS4 → the 118K compendium extraction (random subset, seed = 42) | **411,318** | Computational subset for manuscript-scope training |
| 4 | ARCHS4 → UBERON tissue mapping (keyword + manual curation) | **212,412** | Tissue-labelled subset; un-mappable annotations excluded |
| 5 | + TCGA (bulk tumour samples, GDC release) | **+ 9,400** | Candidate pool gains primary tumour expression |
| 6 | + GTEx (bulk normal tissue, v8 release) | **+ 14,768** | Candidate pool gains normal tissue baseline |
| 7 | + DepMap (cell-line, carried from earlier compendium) | **+ 1,451** | Cell-line samples; removed by cell-line filter (step 9) |
| 8 | Combined candidate pool | **238,031** | Four sources, pre-curation (10 unmappable rows dropped → 238,021) |
| 9 | Cell-line filter: ENCODE/CCLE identifier exclusion | **188,790** | Cell-line samples (DepMap + ARCHS4-flagged) removed |
| 10 | Class balancing: cap of 10,000 samples per UBERON tissue | **146,537** | Dominant tissues capped; final set 123,127 ARCHS4 / 14,072 GTEx / 9,338 TCGA |
| 11 | Train / test counts (85/15 split applied at pool stage) | **118,263 / 28,274** | Used for VAE training and held-out evaluation |

The compendium spans well-represented tissues — blood (n = 8,377 test samples), stem cell (n = 2,189), skin (n = 2,160), brain (n = 1,718), breast (n = 1,542) — and rare categories such as biliary tract (n = 4) and pleura (n = 12). The cell-line exclusion and the per-tissue cap have two effects worth noting at this stage: identifier-matched cell-line samples (all DepMap, plus ARCHS4 entries with cell-line identifiers) are removed, and the cap affects only blood, by far the most over-represented tissue (∼55,000 samples), which is reduced to 10,000; every other tissue falls below the 10,000 threshold and is left unchanged (e.g. lung 9,191, lymphoid 7,280, bone marrow 4,464 training samples). The cap therefore prevents the single dominant class from overwhelming training without altering the smaller classes.

### The latent space is primarily structured by tissue identity, with source effects remaining secondary but detectable

We trained a supervised VAE on the 118,263-sample training split across 16,115 genes (163 near-zero-variance genes were removed). The architecture is a single-hidden-layer encoder feeding a 121-dimensional latent space, trained to simultaneously reconstruct the input expression profile and classify the UBERON tissue label. Hyperparameters were carried over from the 75,619-sample run of the same architecture (see Methods), so that performance differences across compendium sizes reflect data rather than model choices.

The 121-dimensional embeddings were visualized by t-SNE using the Kobak–Berens protocol (Figure 1a). Samples cluster by organ system. Brain forms a tight, isolated neighbourhood; blood and immune tissues form a single large cluster; skin and connective tissue, reproductive organs, GI tract, urinary, and thoracic systems occupy distinct regions. Within broad systems, individual tissues form sub-clusters (e.g. liver vs colon vs pancreas within the GI tract). When the same plot is coloured by data source (Figure 1b), ARCHS4, GTEx, and TCGA samples co-cluster within their respective tissue neighbourhoods rather than forming source-specific islands. On held-out data (n = 28,274 labelled samples), the classifier reaches 94.9% balanced accuracy and 96.2% weighted F1 score across the 42 tissue categories represented in the test split (Figure 2; Supplementary Figure S1). Accuracy is highest for tissues with distinctive transcriptomes — pituitary, pleura, salivary gland, spinal cord, and vagina at 100%; prostate, muscle, breast, and blood vessels above 98%. The lowest-scoring classes are those with very few test samples (biliary tract n = 4, eye n = 24, thymus n = 32, spleen n = 68), and all remain above 75%. The lymphoid and bone marrow classes reach 91.9% and 95.5% (Table 2). Capping the dominant blood class helps the model resolve the related haematopoietic tissues rather than defaulting to blood. Per-class accuracy and sample counts are shown in Figure 2; the normalized confusion matrix is in Supplementary Figure S1.

**Table 2.** Data scaling and curation across three training runs of this study. Per-sample reconstruction quality is the metric most responsive to data scale and curation; per-gene fidelity and classification accuracy continue to gain but approach saturation. The 38K and 75K runs are scaling reference points; the 118K run is the primary model reported here.

| Metric | 38K-sample run | 75K-sample run | 118K-sample run |
| --- | --- | --- | --- |
| Cell lines excluded | No | No | Yes |
| Class balanced | No | No | Yes (10K/tissue cap) |
| **Per-gene $\rho$ (Denoising)** | 0.892 | 0.919 | 0.934 |
| **Per-sample $\rho$ (Denoising)** | 0.642 | 0.773 | 0.830 |
| **Imputation @ 30% (Denoising)** | 0.825 | 0.855 | 0.879 |
| **Classification BA (Standard)** | 89.5% | 90.7% | 94.9% |
| **Weighted F1 (Standard)** | 92.4% | 93.7% | 96.2% |
| **TARGET overall accuracy** | — | 86.6% | 84.6% |
| **Peak RAM during data loading** | ~30 GB CSV | ~50 GB CSV | 25 GB HDF5 |

$\Delta$ per-sample $\rho$ : 38K → 75K: +27%; 75K → 118K: +7.4%; total 38K → 118K: +38%.

**Figure 1.**
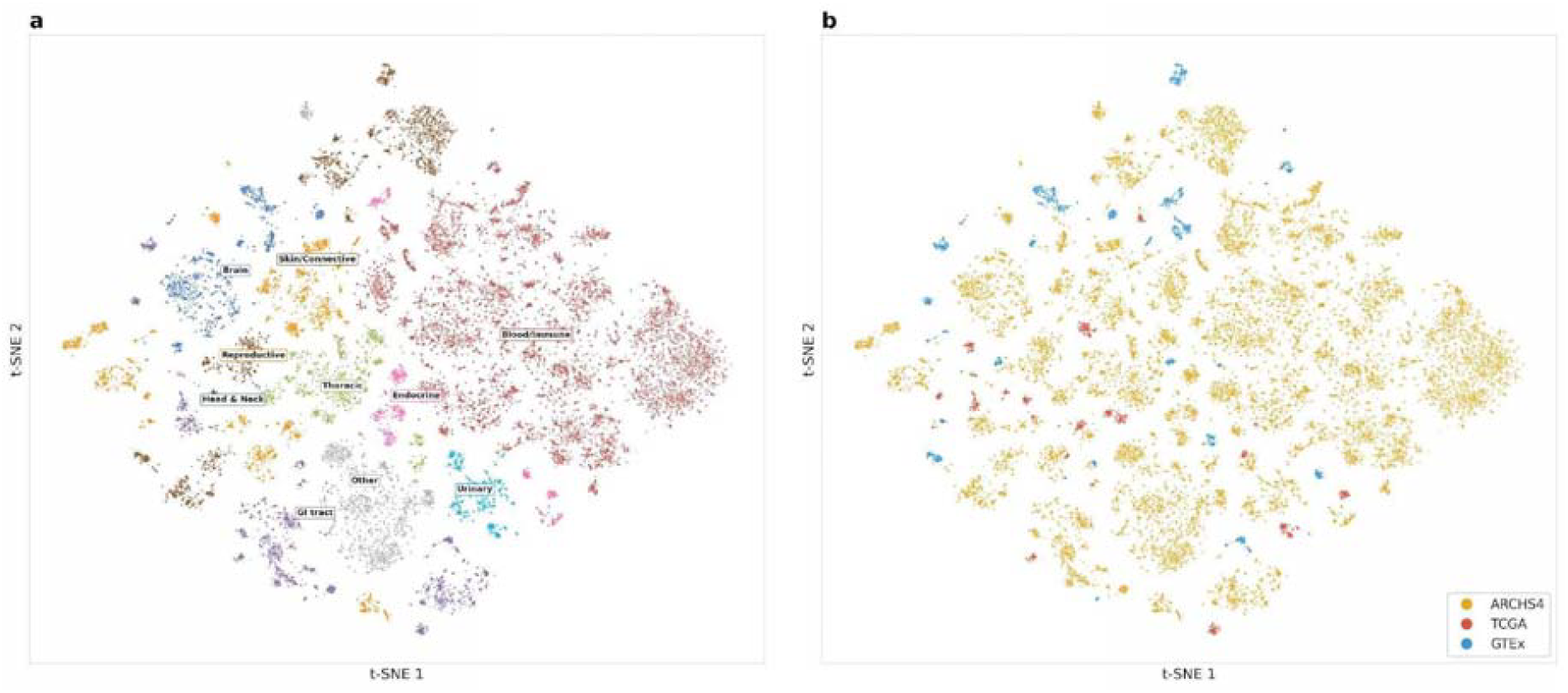
The 118K latent space is primarily structured by tissue identity, with source effects secondary. t-SNE projection of the 121-dimensional latent space for held-out samples (n = 28,274), using the Kobak–Berens protocol (PCA initialisation, perplexity 30, learning rate n/12, cosine metric). (a) Coloured by organ-system grouping of the 42 UBERON tissue categories. Brain, Skin/Connective, Blood/Immune, Reproductive, Thoracic, GI tract, Urinary, Endocrine, Head & Neck, and Other systems occupy distinct regions; within each region, individual tissues form sub-clusters. (b) The same embedding coloured by data source (ARCHS4, TCGA, GTEx). Samples are organised primarily by tissue rather than by source; quantitative source-mixing metrics for this embedding (LISI 1.02/3.0; kNN source mixing 0.012) are interpreted in the main text

**Figure 2.**
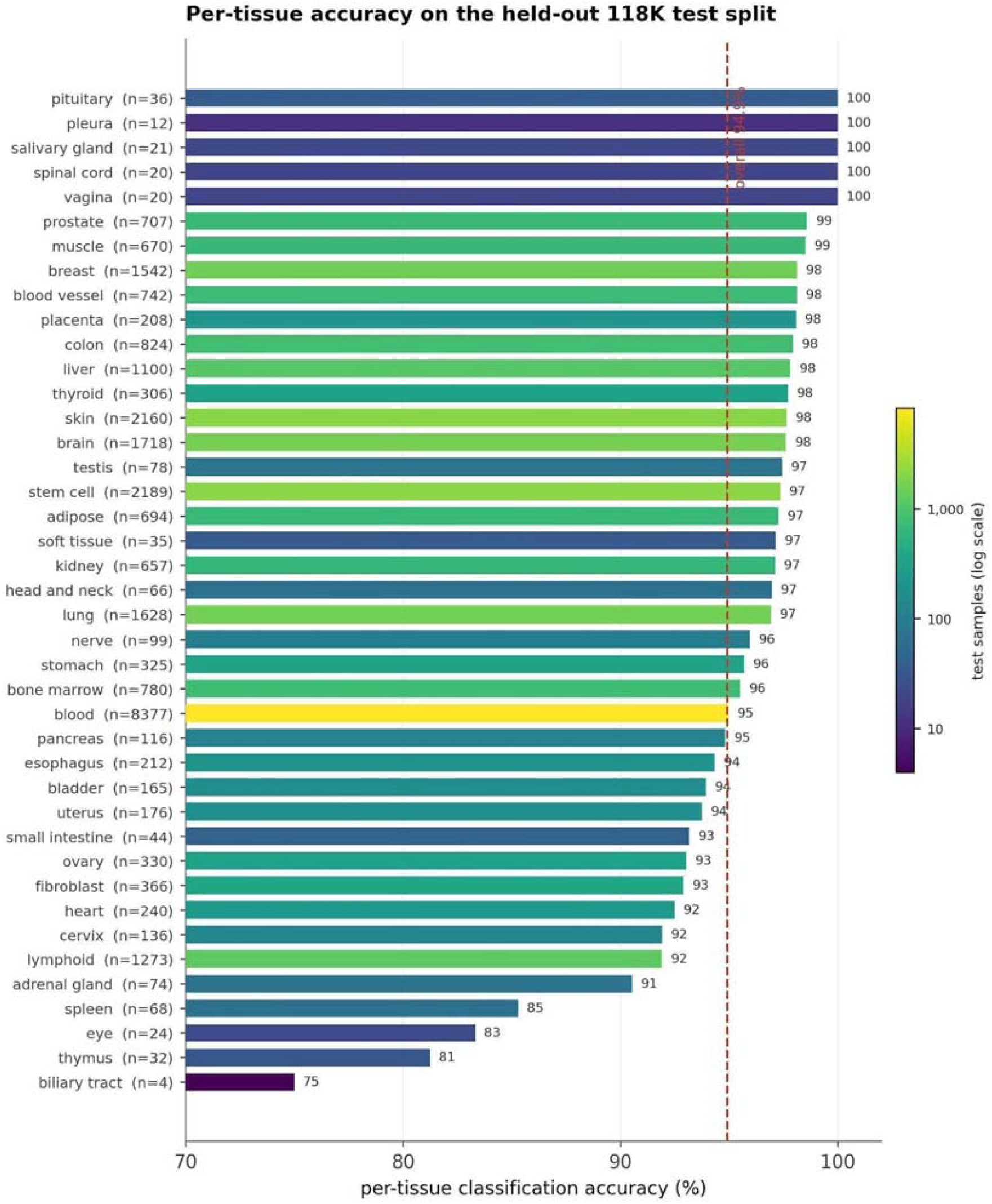
Per-tissue classification accuracy on the held-out 118K test split. Balanced accuracy per UBERON tissue category, sorted from highest (top) to lowest (bottom). Bars are coloured by held-out test sample size (log scale; colour bar at right), and sample sizes (n) are shown beside each tissue label. The dashed vertical line marks the overall balanced accuracy (94.9%; weighted F1 96.2%). Accuracy is uniformly high across well-sampled tissues and degrades only for the rarest categories: the four lowest-accuracy tissues (biliary tract 75%, n = 4; thymus 81%, n = 32; eye 83%, n = 24; spleen 85%, n = 68) are all small classes for which test-set estimates are noisy, whereas every tissue with more than ∼100 test samples exceeds 90%. Previously low-count classes such as lymphoid (91.9%, n = 1,273) and bone marrow (95.5%, n = 780) recover after class-balanced training.

Quantitatively, source effects remained secondary to tissue identity in the latent space. kNN source mixing (k = 20) was 0.012, indicating that only 1.2% of a sample’s nearest neighbours originated from a different source. This represents a further reduction relative to the 75K-sample model (0.047), consistent with the exclusion of cell-line samples and improved class balancing in the curated 118K compendium (Table 2). Since source mixing is measured within tissue clusters, that residual 1.2% means neighbours sharing a sample’s tissue but from a different source — not samples grouping by source (Figure 1). Full mixing would not necessarily be desirable either: the sources differ in tissue composition (GTEx mostly normal tissue, TCGA mostly tumours), so some source structure is expected. We did not test whether the residual within-tissue structure reflects disease state (tumour vs normal) or residual source differences, and the two cannot be separated from the present analysis. The source signal is still detectable, but tissue identity dominates the geometry.

### Reconstruction and gene-imputation benchmarking

Beyond classification, we evaluated how faithfully the VAE reconstructed the original expression profile of held-out samples (Figure 3). We compared two model variants trained with identical architecture and hyperparameters, differing only in whether 20% of input genes were randomly zero-masked during training (Denoising) or not (Standard) (see Methods). Per-gene Spearman correlation between original and reconstructed expression was similar for the two variants (median ρ = 0.935 Standard, 0.934 Denoising; Figure 3a, Supplementary Figure S2). Per-sample reconstruction quality was also close (median ρ = 0.829 Standard, 0.830 Denoising; Figure 3b). The bimodal shape of the per-sample distribution — a dominant peak around ρ ≈ 0.85 with a longer left tail — separates tissues with strong, distinctive expression programmes from those with more generic profiles. Across the three training runs of this study, per-sample ρ rose at each step (0.604 → 0.773 → 0.830 across 38K, 75K, and 118K samples), and per-gene ρ showed the same trend (0.892 → 0.919 → 0.935) (Table 2).

**Figure 3.**
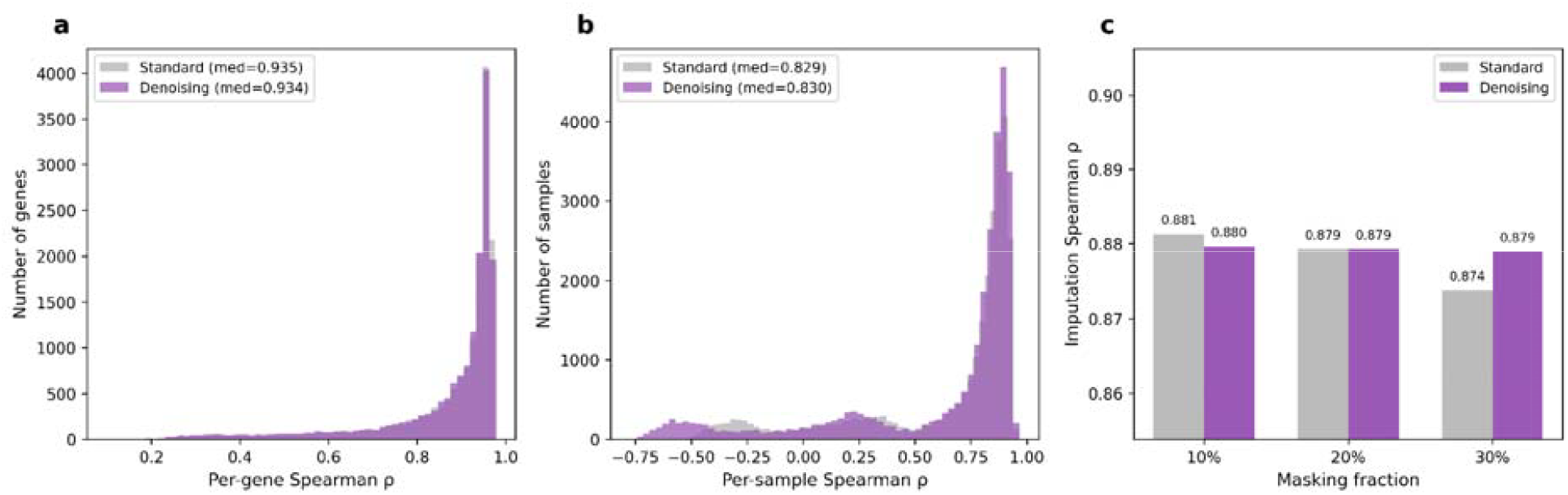
Reconstruction and imputation performance of Standard versus Denoising variants. (a) Per-gene Spearman correlation between original and reconstructed expression, across 16,115 genes on the held-out test split. Median ρ = 0.935 (Standard) vs 0.934 (Denoising). (b) Per-sample Spearman correlation across 28,274 held-out samples. Median ρ = 0.829 (Standard) vs 0.830 (Denoising). The bimodal distribution separates tissues with distinctive expression programmes (right peak) from those with more generic profiles (left tail). (c) Imputation Spearman ρ between zero-masked and reconstructed expression, across three masking fractions. Imputation accuracy is comparable for the two variants across masking fractions (within 0.005; 30% masking: 0.874 Standard, 0.879 Denoising on the full held-out test set)

Gene imputation under random masking gives comparable results for the two variants (Figure 3c). Across masking fractions the two are within 0.005 of each other on the full held-out test set (10%: 0.881 Standard vs 0.880 Denoising; 20%: 0.879 vs 0.879; 30%: 0.874 vs 0.879) — within the variation expected between training runs of the same model. We therefore treat the variants as equivalent; the denoising objective is retained as a no-cost default that may help when downstream inputs are incomplete or cross-platform.

### Model validation on held out dataset: TARGET paediatric cancers

We validated the model on an independent cohort of 734 paediatric tumour samples from TARGET (15,16), achieving 84.6% agreement with the expected tissue of origin. TARGET is an independent paediatric oncology programme. Although our compendium draws on ARCHS4, no TARGET samples were present in our extracted training data (verified by sample-identifier screening), so the cohort was unseen during training. Because no paediatric tumours or developmental-biology annotations were included during training, transfer to this cohort is an out-of-distribution test.

Many paediatric cancers arise from cells that, during development, give rise to specific adult tissues (17-19); the question is whether representations learned from adult bulk transcriptomes capture enough of this developmental relationship to recover the expected tissue of origin. TARGET expression profiles were aligned to the 16,115 model genes, per-gene z-score harmonized to training distributional statistics, and encoded into the 121-dimensional latent space. Each TARGET sample was then classified by k-nearest-neighbour (k = 5, Euclidean) against the 118K reference embeddings (Figure 4). Expected tissue assignments were derived from established developmental biology and defined a priori for each cancer type (Supplementary Table S2). Leukaemias arise from haematopoietic progenitors, so their expected adult tissues are blood, bone marrow, and lymphoid tissue. Neuroblastoma arises from the neural crest, with brain, nerve, and adrenal gland as the closest adult correlates. Wilms tumour and rhabdoid tumour of the kidney both arise from the intermediate mesoderm, which gives rise to kidney, ovary, uterus, and testis in the adult. Overall, the model assigned the expected tissue of origin for 84.6% of samples (621/734) (Figure 4).

**Figure 4.**
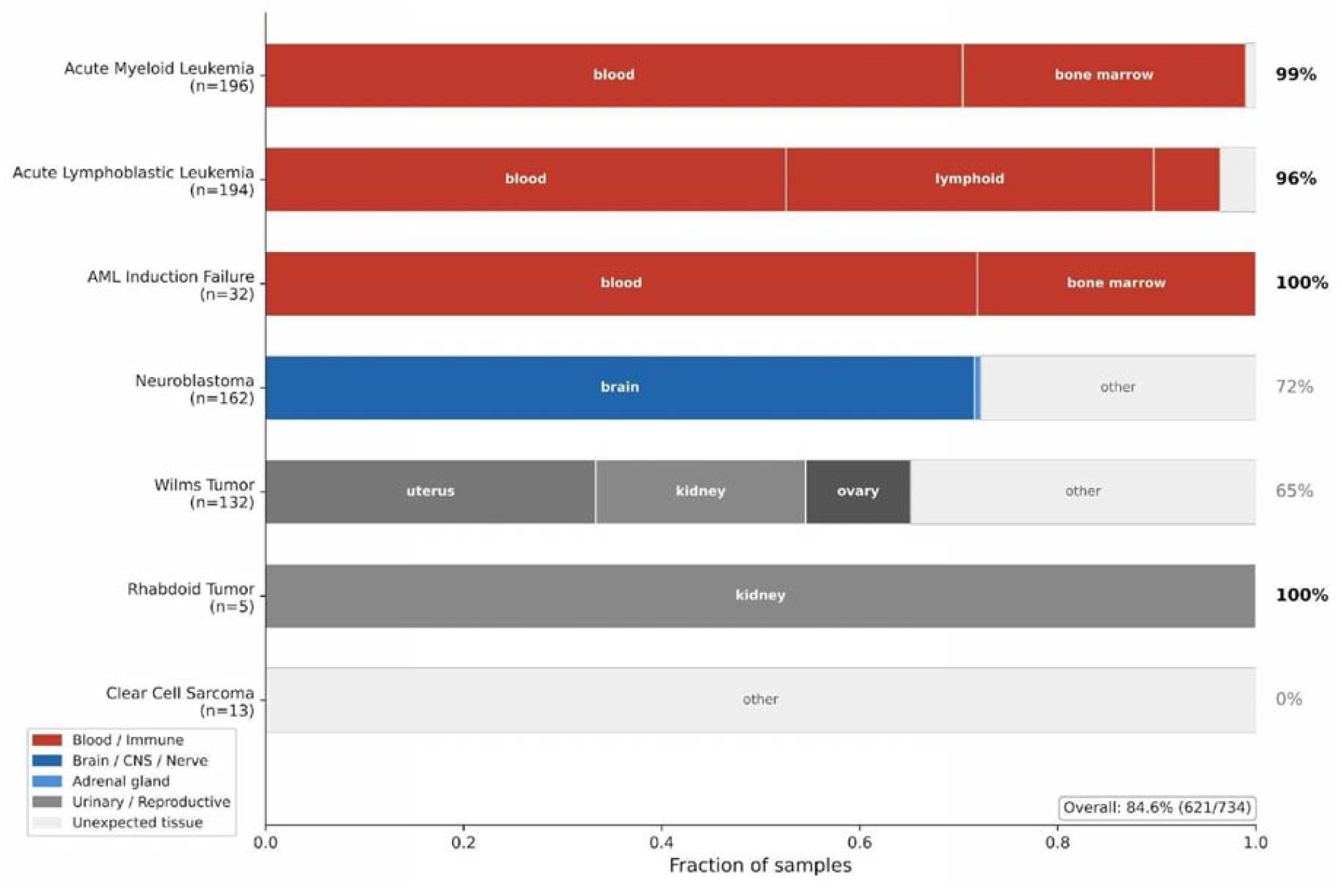
Transfer to TARGET paediatric cancers. Each row is one TARGET cancer category (n = 734 samples across seven categories); horizontal bars show the fraction of samples whose nearest neighbour in the 121-dimensional latent space falls in each expected adult tissue category. Expected tissues are color-coded by developmental origin: Blood/Immune (haematopoietic), Brain/CNS/Nerve and Adrenal gland (neural-crest), Urinary/Reproductive (intermediate-mesoderm), and Unexpected tissue (light grey). Right-side percentages report the fraction of expected-tissue assignments per cancer type; overall accuracy 84.6% (621/734). Clear Cell Sarcoma of the kidney (BCOR-driven, no adult-tissue correlate) maps to no expected tissue.

### Leukaemias map to blood and bone marrow

Haematopoietic cancers comprise the largest category the model was tested on. Acute myeloid leukaemia, which arises from myeloid progenitors in the bone marrow (17), maps to blood or bone marrow in 99% of TARGET samples (n = 196). AML induction failure — the molecularly distinct refractory variant — reaches 100% (n = 32). Acute lymphoblastic leukaemia, which originates from lymphoid progenitors, maps to blood or lymphoid tissue in 96% of cases (n = 194); the corresponding rate from the 75K-sample training run was 65% (Table 2). The difference tracks the per-tissue cap introduced in the 118K compendium, which prevents the lymphoid category from being absorbed into the dominant blood signal during training. Rhabdoid tumour of the kidney, a non-haematopoietic control, maps to the kidney in 5 out of 5 samples (Figure 4).

### Neuroblastoma maps predominantly to brain

Neuroblastoma arises from neural-crest cells of the developing sympathetic nervous system (18). Neural crest is not a category in the 42-tissue UBERON ontology used here; the closest adult anatomical correlate is brain. In TARGET, 72% of neuroblastoma samples (n = 162) map to brain. The remaining 28% distribute across categories without a clear developmental rationale. The 75K-sample training run produced a tighter 99% mapping to brain plus adrenal gland, but that earlier figure was possibly inflated as it was carried largely by neural-crest-like cell-line samples, which the 118K compendium excludes by design. The 72% therefore represents a more conservative estimate of transfer from a representation free of cell lines (Figure 4).

### Wilms tumour distributes across intermediate-mesoderm-derived tissues

Wilms tumour arises from persistent nephrogenic rests — embryonic remnants of the intermediate mesoderm, which also gives rise to the kidney, the gonadal ridge, and the Müllerian and Wolffian ducts (precursors of uterus, ovary, and testis) (19). Wilms tumour samples in TARGET (n = 132) distribute across these mesoderm-derived adult tissues: uterus (33%), kidney (21%), ovary (11%) — 65% combined. The remaining 35% sit largely within other urogenital and reproductive categories. Wilms tumour samples do not concentrate on a single tissue label; they occupy a region of latent space defined by multiple intermediate-mesoderm-derived adult tissues (Figure 4).

### Clear cell sarcoma of the kidney: no match

Clear cell sarcoma of the kidney (n = 13) maps to no expected tissue (0% accuracy). The tumour is driven by BCOR internal tandem duplications (20) and has no transcriptomic correlate among the adult tissues represented in our training dataset. The result suggests that paediatric tumours driven by tumour-specific genomic markers are not recoverable from the transcriptome representations of adult tissues (Figure 4).

### Comparison with baseline and zero-shot approaches

We compared the supervised VAE against two classes of alternatives on the same held-out 118K test split: (i) kNN classification applied directly to raw or reduced expression spaces, and (ii) zero-shot classification using a pre-trained single-cell foundation model.

Five kNN-based baselines were evaluated, spanning representation choices from raw 16,115-gene expression to several dimensionality-reduction pipelines (Figure 5a; Methods). The strongest kNN configuration, top-2,000 highly variable genes with k = 5 cosine kNN, reached 93.4% balanced accuracy. The all-gene kNN classifier reached 92.6%, and PCA-and UMAP-based reductions ranged from 84.2% to 92.0%. The supervised VAE reached 94.9% in a 121-dimensional latent space. The value of the supervised VAE is not merely this improvement in classification accuracy, but the representation it produces: a 121-dimensional embedding (135-fold compression) that transfers to unseen cohorts and supports reconstruction and imputation, which are tasks that a kNN classifier built on raw expression space cannot perform.

**Figure 5.**
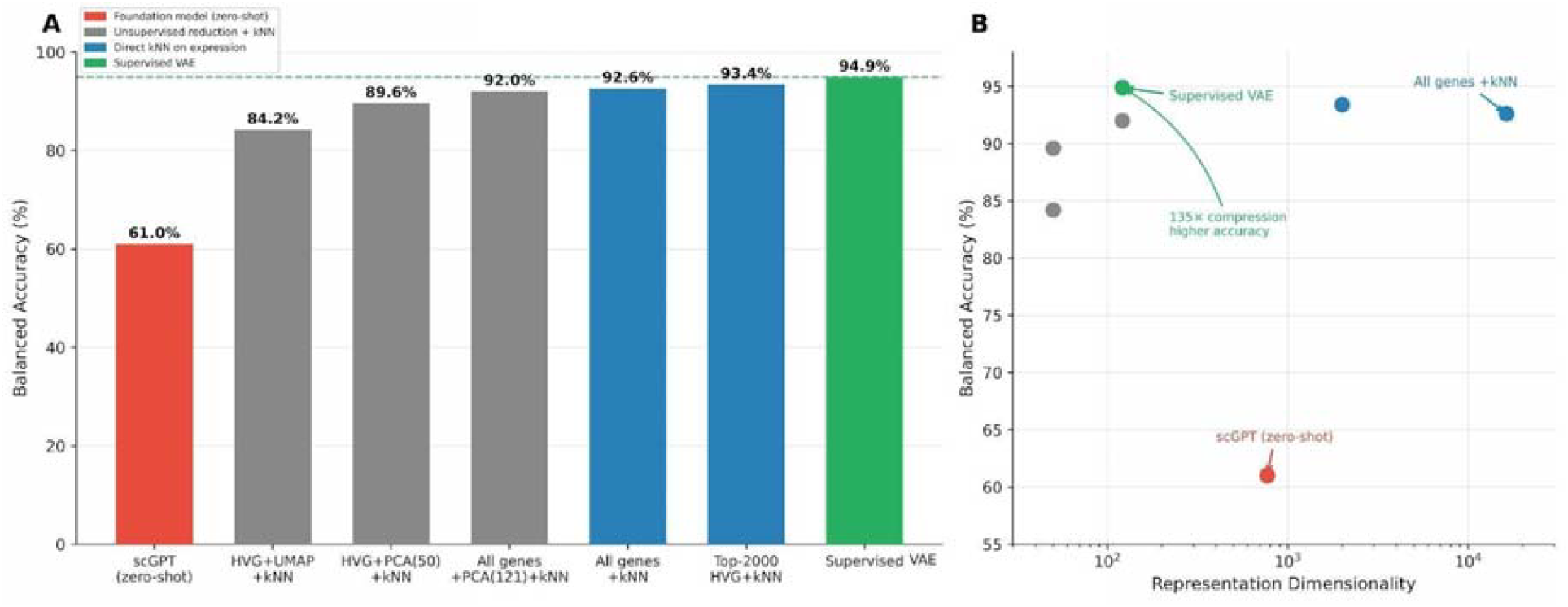
The supervised VAE outperforms kNN baselines and a zero-shot single-cell foundation model. (a) Balanced accuracy on the held-out 118K test split, comparing scGPT in zero-shot mode (red); four unsupervised reduction + kNN baselines (grey: HVG+UMAP, HVG+PCA(50), all genes + PCA(121), top-2,000 HVG); two direct kNN-on-expression baselines (blue); and the supervised VAE (green). Dashed green line marks the supervised VAE accuracy (94.9%). (b) Accuracy–dimensionality trade-off for the same set of methods. The supervised VAE achieves the highest accuracy at 121 dimensions — a 135-fold compression relative to the all-genes kNN representation. scGPT (zero-shot) sits well below at 768 dimensions, reflecting the domain gap between single-cell pre-training and bulk RNA-seq evaluation.

scGPT, applied in zero-shot mode to bulk RNA-seq inputs without fine-tuning, reached 61.0%. The reduced performance of scGPT can be explained by the fact thatscGPT is trained on single-cell transcriptomes, while our test set is bulk RNA-seq. Zero-shot transfer from single-cell to bulk rna-seq recovers tissue identity well above chance, but well below specialized models.

Beyond accuracy, the methods also differ in the dimensionality of the representations they produce. The accuracy–dimensionality trade-off is shown in Figure 5b. The supervised VAE reaches the highest accuracy at the lowest dimensionality (135-fold compression relative to all-gene kNN), while remaining a small enough model to train and run on a single GPU.

We also compared the supervised VAE against BulkFormer, a recent bulk-RNA-seq-specific foundation model (12). Applied out of the box, using the authors’ default sample-level feature extraction without task-specific fine-tuning, BulkFormer was evaluated on the TARGET cohort with the same k-nearest-neighbour protocol and developmental-expectation rubric used for our model (Figure 6). The two models agreed on the haematopoietic cancers (both above 97%) and were close on neuroblastoma (72.2% versus 75.9%). Overall, though, the supervised VAE found the expected tissue of origin more often (84.6% versus 76.8%). Most of the gap was Wilms tumour: our VAE put 65.2% of these samples in intermediate-mesoderm tissues, while BulkFormer reached only 13.6% and assigned most of them to brain, which is a different germ layer (Figure 6b). Per-category accuracies for both models are summarized in Supplementary Table S3. It is important to note that BulkFormer was pretrained on GEO and ARCHS4 and has almost certainly seen TARGET samples, whereas these samples were never included in our model training. So the transfer is genuinely out-of-distribution only for our model.

**Figure 6.**
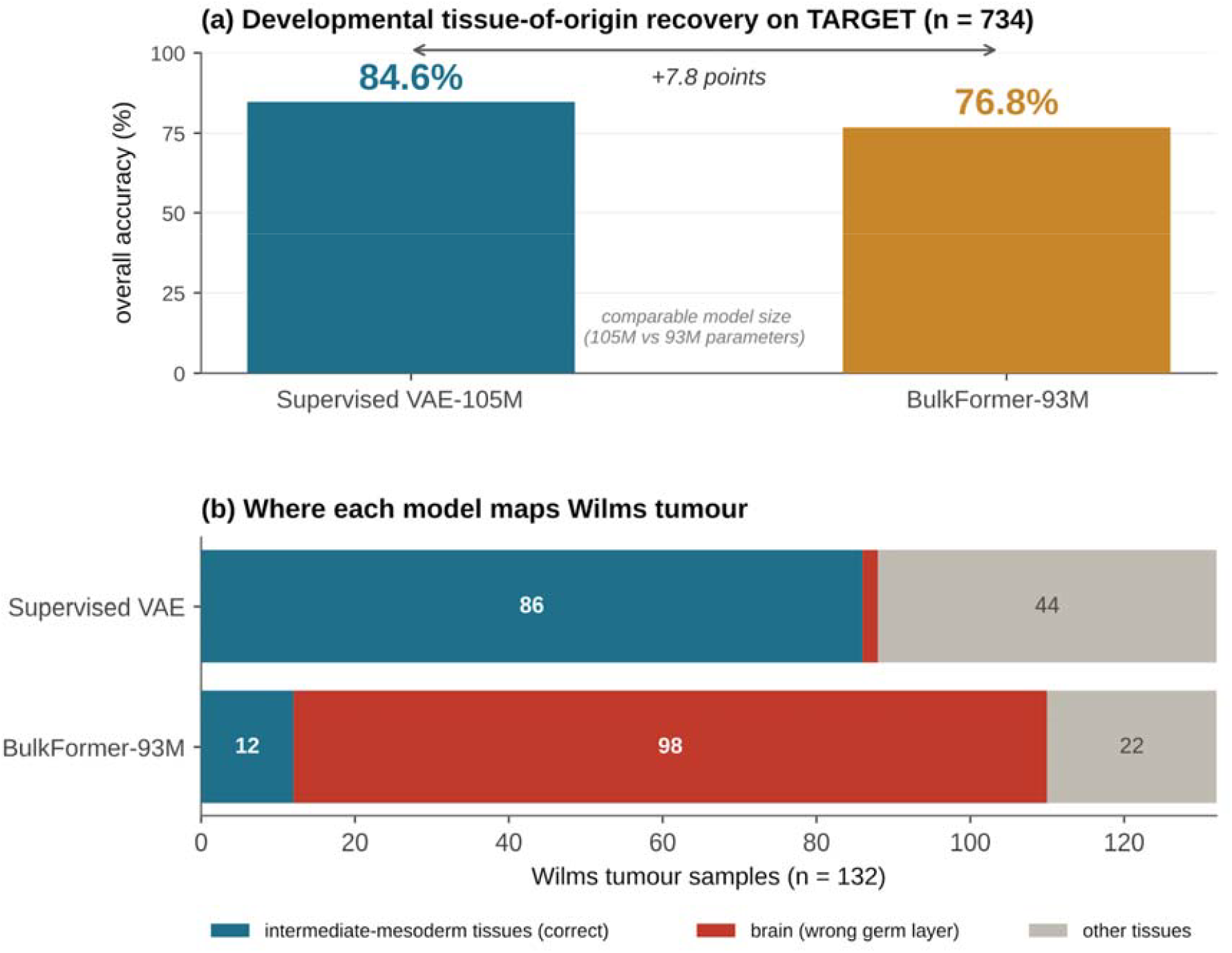
The supervised VAE recovers developmental tissue-of-origin relationships more accurately than an out-of-the-box bulk RNA-seq foundation model. Comparison of the supervised VAE (this work; ∼105M parameters) with BulkFormer-93M, a bulk-transcriptome foundation model of comparable size, on the 734-sample TARGET paediatric cohort. Both models were embedded and classified by k-nearest-neighbour (k = 5, Euclidean) against the held-out reference and scored against the same developmental-expectation rubric; each model received its native input representation, and BulkFormer was applied out of the box without task-specific fine-tuning. (a) Overall developmental tissue-of-origin accuracy: the supervised VAE attains 84.6% versus 76.8% for BulkFormer. (b) Predicted tissue assignments for the 132 Wilms tumour samples, the category driving the overall difference. Intermediate-mesoderm-derived tissues (kidney, uterus, ovary; correct origin) are shown in teal; brain, an ectoderm-derived tissue representing a germ-layer error, in red; all other tissues in grey. The VAE assigns 86 of 132 samples to mesoderm-derived tissues, whereas BulkFormer assigns 98 of 132 to brain.

### Effects of data scaling and curation

The trajectory from 38,296 → 75,619 → 118,263 samples shows consistent improvement across reconstruction and classification metrics (Table 2). The 75K → 118K transition is not purely a comparison of dataset scales, however, because the 118K dataset also excludes cell lines and caps the representation of tissues at a maximum of 10,000 samples per category. The two changes act in the same direction: per-tissue capping raises performance on previously under-represented classes (lymphoid +19, bone marrow +16 percentage points), while cell-line exclusion lowers the source-mixing metrics (LISI 1.09/4.0 → 1.02/3.0; kNN source mixing 0.047 → 0.012), indicating stronger source separation rather than better integration — a consequence of removing the single-source cell-line cluster, as discussed above; a full metric-by-metric comparison of the two versions is given in Supplementary Table S1. On the other hand, while the latent space learned in the 118K-sample run is more broadly biologically organized, the 75K-sample run, which included the cell line samples, displayed better performance on specific cases, such as Neuroblastoma/Wilms tumors, because they were informative on rare developmental signatures..

With respect to the transcriptome reconstruction and missing data imputation tasks, the improvements across the three runs diminish with increasing sample sizes: per-sample Spearman ρ rises by 27% from 38K to 75K, and by 7.4% from 75K to 118K (per-gene Spearman shows the same pattern, Δ = 3.0% and 1.6%; Table 2). The 118K compendium changes what the representation captures, not just how well. The per-tissue cap lifts acute lymphoblastic leukaemia transfer from 65% to 96% (Table 2), recovering a developmental signal that the blood-dominated 75K set had absorbed. So scaling here improves coverage of under-sampled lineages. The main tissue-level variance is already captured at 118K. Scaling further to the 605K-sample bulk corpus should therefore improve coverage rather than the metrics themselves — reaching rarer tissues, finer cell-state variation within well-represented tissues, and denser reference embeddings for nearest-neighbour transfer. We do not expect it to significantly raise tissue-classification or reconstruction scores.

## Discussion

A supervised VAE trained on the 118,263-sample tissue-curated, class-balanced dataset learns tissue representations that classify a held-out split at 94.9% balanced accuracy, transfer to a paediatric cancer cohort absent from training, and reflect the developmental signatures that connects paediatric tumours to their adult tissues of origin. We discuss the biological, data-scaling, and methodological implications of these results in turn, and then their limitations and natural extensions.

When applied to 734 TARGET paediatric tumours absent from training, the model assigned the developmentally expected tissue of origin for 84.6% of samples, compared with 86.6% in the 75K-sample run. As reported above, the per-cancer mapping recovers the expected developmental lineage in most categories, and the per-tissue cap is what lifts the weakest 75K class (acute lymphoblastic leukaemia) into line with the rest. We tested the cost of dropping cell lines directly. Keeping the ARCHS4 cell lines under the same capping policy and the same random seed, so that cell-line inclusion was the only thing that changed (Supplementary Figure S3), overall TARGET accuracy was 87.1% with cell lines and 84.6% without. The whole difference was in neuroblastoma (81% versus 72%); every other cancer type was unchanged, and Wilms tumour stayed within two points (63% versus 65%). So removing cell lines costs little, and only in the one lineage—neural crest—that cell lines represent best.

Our model displayed better performance than BulkFormer, however, task-specific fine-tuning of BulkFormer might narrow this gap; the comparison indicates that a tissue-supervised representation captures the developmentally subtle intermediate-mesoderm mapping that an out-of-the-box bulk foundation model does not.

The supervised VAE reaches 94.9% balanced accuracy in a 121-dimensional space; the strongest kNN baseline (top-2,000 highly variable genes) reaches 93.4%. In absolute terms the VAE margin is small (+1.5 percentage points), but the representation it produces is 135-fold compressed, transfers to unseen cohorts, and supports both reconstruction and imputation — properties that nearest-neighbour lookup in raw expression space does not provide.

The latent space is primarily structured by tissue identity, with source effects remaining secondary but detectable, although no explicit batch correction (ComBat, Harmony, adversarial training) was applied. The supervised tissue-classification objective is sufficient in this setting to absorb source variation into the tissue axis while preserving real biological differences (tumour vs normal) within each tissue. Stronger batch correction may help applications that require it, and adversarial or contrastive objectives could be added; the supervised approach used here avoids needing to define what constitutes a “batch” in advance.

The denoising variant produces near-identical per-gene and per-sample reconstruction and imputation to the standard model; the two are equivalent in performance. The denoising objective adds no computational cost and no architectural change, so it is a reasonable default when downstream tasks involve incomplete or cross-platform inputs. One limitation of the current implementation is the use of zero-substitution rather than true masked-autoencoder (MAE) training: the MLP encoder operates on a fixed-length input vector, so masked positions cannot be excluded from the encoder computation. A transformer encoder would enable proper MAE training and may strengthen the denoising advantage.

Several directions follow naturally from this work. The cell-line ablation reported above isolates one of the two curation axes; a complementary control that varies only the per-tissue cap (balanced versus unbalanced at fixed composition) would similarly isolate the contribution of class balancing from that of sample count. A normalization layer learned within the model would replace the heuristic per-gene z-score alignment currently used for cross-platform transfer. Because the Flexynesis model tolerates missing labels, the unlabelled samples set aside here could also be co-trained on the reconstruction objective; they would not contribute to the classification loss but could strengthen the reconstruction and imputation arm of the model. Finally, a full run on the 605,614-sample bulk-filtered ARCHS4 corpus is the natural extension of this work, expected to improve coverage of rare tissues and the density of the reference embeddings rather than improving the model performance metrics for classification and transcriptome reconstruction

The 42-category UBERON ontology is a compact representation of anatomy, which doesn’t comprise fine-grained tissue type (and subtype) labels. For instance, “brain” includes cerebellum and cortex, which have very different transcriptomes; or “kidney” does not distinguish cortex from medulla. The cross-platform normalization used for TARGET (per-gene z-score alignment to training statistics) is a heuristic; replacing the fixed per-gene z-score normalization with a trainable normalization layer — learned during normal training, not fine-tuned on held-out data — is a direction for future work.

The model evaluations performed in this study are restricted to tissue label classification, transcriptome reconstruction, and missing-data imputation. Downstream uses such as survival prediction, drug response, and subtype discovery were not tested, as these require paired clinical and outcome labels that the public expression compendia used here do not provide.

## Materials and Methods

### Data assembly and tissue curation

RNA-seq expression data were obtained from three public resources: TCGA (via the Genomic Data Commons; n = 9,400 tumour samples), GTEx (v8 release; n = 14,768 normal tissues), and ARCHS4 (uniformly processed GEO submissions) (2,3,5). ARCHS4 v2.5 contains 888,821 human RNA-seq samples drawn from diverse experimental contexts, including a mixture of bulk and single-cell submissions. We filtered the samples to keep only bulk transcriptome samples using the built-in single-cell probability score (excluding samples with score ≥ 0.5), yielding 605,614 bulk RNA-seq samples. From this pool, 411,318 samples were extracted and their free-text tissue annotations mapped to UBERON categories (21) by keyword matching followed by manual curation, yielding 212,412 tissue-labelled samples.

The candidate pool of 236,580 samples (212,412 ARCHS4 + 14,768 GTEx + 9,400 TCGA) was reduced first by removing 10 samples with unmappable tissue annotations (236,570), then by two filters: (i) ARCHS4 cell-line-flagged samples were excluded by pattern matching against ENCODE/CCLE-style identifiers (188,790 remaining); because this filter is identifier-based, a residual number of ARCHS4 samples annotated as cell lines only in free-text descriptions were not caught and remain, labelled by their tissue of origin, and (ii) each UBERON tissue category was capped at 10,000 samples. The final compendium contains 118,263 training samples and 28,274 test samples (146,537 total; 123,127 ARCHS4, 14,072 GTEx, 9,338 TCGA) across 42 UBERON tissue categories.

Expression values were log2-transformed (TPM or RPKM depending on source). Gene identifiers were harmonized to HGNC symbols (22), yielding 16,292 shared genes. After removing 163 genes with near-zero variance, 16,115 genes were retained for training. Expression values were standardized using sklearn StandardScaler (23) fitted on the training split only. Samples without a UBERON tissue annotation were excluded from supervised training.

### Training data assembly across three scales

Three training compendia were assembled during the development of this work, and results across all three are reported in Table 2 and discussed in the Results.

The 38,296-sample compendium was an initial experiment drawn from TCGA, GTEx, DepMap, and a 20,000-sample subset of ARCHS4, mapped to 43 UBERON tissue categories. Cell-line samples were retained, and tissue categories were not class-balanced.

The 75,619-sample compendium extended the ARCHS4 contribution to approximately 50,000 samples (training/test split: n = 64,279 / 11,340), keeping TCGA, GTEx, and DepMap as before, across the same 43 UBERON tissue categories. Cell-line samples were retained, and tissue categories were not class-balanced; low-count classes such as lymphoid (n = 179 test samples) and bone marrow (n = 262) were under-represented relative to dominant categories such as blood (n = 1,266).

In order to address the issue with under/over representation of tissue types, we curated a 118K-sample dataset. The 118,263-sample compendium is the curated compendium used as the primary training set in this work, described in the section above. It differs from the 75,619-sample compendium in two ways: cell-line samples are excluded, and tissue categories are size-capped at 10,000 samples each. The 38K and 75K runs are reported here as scaling reference points; the 118K run is the primary model.

The same VAE architecture and the same Bayesian-optimized hyperparameters were used across all three runs, so differences between them reflect training-data changes rather than model choices.

### HDF5-backed dataloader (H5DataImporter)

Loading the 118,263 × 16,115-gene matrix via the conventional pandas-CSV pipeline peaks above 60 GB of RAM, because pandas reads CSV values as float64 by default. We implemented H5DataImporter, a subclass of the Flexynesis DataImporter (21) that overrides only the read_data() and validate_data_folders() methods. The new path loads expression matrices from HDF5 (24) as native float32 arrays and returns the DataFrame that downstream Flexynesis components expect, with no other modification. Peak memory during data loading drops to approximately 25 GB for the 118K × 16K matrix.

CSV input files are converted to HDF5 once using a chunked-write converter (csv_to_h5.py) that pre-allocates the target array and writes samples-as-rows blocks. All other Flexynesis components (cleanup, NaN imputation, label encoding, scaling, torch dataset construction) operate unchanged on the HDF5-loaded DataFrame.

### Model architecture and training

The supervised VAE used in this work has: (i) a single-hidden-layer encoder (16,115 → 3,226 → 121 dimensions), The latent dimension (121), hidden-dimension factor (0.20), learning rate, and batch size were selected by Bayesian hyperparameter optimization on an initial training run and held fixed for all training-set sizes and for the denoising variant. (ii) a symmetric decoder, and (iii) a two-layer classification head (121 → 32 → 42 classes). The encoder outputs mean and log-variance vectors for the approximate posterior; latent codes are sampled via reparameterisation (25). The loss sums MMD regularisation (26), MSE reconstruction, and cross-entropy classification.

Training used Adam (lr = 1.72 × 10^−3^, batch size 32) (27) with early stopping on validation loss (patience 10 epochs). On the 118K compendium, the Standard model early-stopped at epoch 37 (val_loss = 0.8012) and the Denoising model at epoch 44 (val_loss = 0.8076). The corresponding values for the 75,619-sample run were epoch 17 (val_loss = 0.9248) for the Standard model. The same architecture and hyperparameters were used for all three training runs (38K, 75K, 118K). Models were trained with: `python train_denoising_vae_h5.py --data_path <compendium> --outdir <results> --mask_fraction 0.2 --epochs 500 --early_stop_patience 10 –also_train_standard.`

Hyperparameters were selected by Bayesian optimization on an initial training run and then held fixed across all three training-set sizes (38K, 75K, 118K) and for the denoising variant, so that performance differences reflect data rather than model choices. The selected configuration was: latent dimension 121, hidden-dimension factor 0.200, supervisor hidden dimension 32, learning rate 1.72 × 10^−3^, and batch size 32. The optimization ran for three iterations, searching latent dimension and batch size: the trials evaluated latent dimension 105 (batch size 32), latent dimension 67 (batch size 128), and latent dimension 121 (batch size 32), and the third minimized validation loss and was selected. Batch size 32 was therefore chosen over 128 by the search rather than retained as a default. Training used the Flexynesis supervised_vae model (equivalent command: flexynesis -- model_class supervised_vae --data_types gex --target_variables uberon_tissue --hpo_iter 3 -- early_stop_patience 10). The standard and denoising variants deliberately share this configuration: because the denoising model is evaluated as a controlled ablation of the standard VAE, holding hyperparameters identical ensures that any difference between them is attributable to the denoising objective alone rather than to separate tuning.

### Denoising variant

During each training step, 20% of input gene expression values are randomly set to zero per sample, and the reconstruction loss is computed against the original unmasked values. Architecture and hyperparameters are identical to the Standard model. At inference, no masking is applied.

### t-SNE visualisation

121-dimensional latent embeddings of the 118K test set (n = 28,274) were projected to two dimensions using the Kobak–Berens protocol (28): PCA initialisation, perplexity 30, learning rate n/12 = 2,356, 750 iterations, cosine metric. Implemented in openTSNE v1.0 (29).

### Source mixing metrics

Source mixing in latent space was assessed by two metrics: (i) kNN source mixing (k = 20) — the fraction of a sample’s nearest neighbours that come from a different data source; (ii) local inverse Simpson index (LISI) (8) — the effective number of sources among neighbours, ranging from 1.0 (single source) to N (equal mixing across N sources). The LISI ceiling for the 118K compendium is 3.0 because DepMap is excluded; for the 75,619-sample four-source run the ceiling was 4.0. The kNN source-mixing metric is independent of the source count and is the more directly comparable statistic across compendia.

### Baseline benchmarks

Five kNN-based baselines were evaluated on the 118K compendium using the same train/test split as the VAE: (i) all 16,115 genes + kNN (k = 5, cosine); (ii) top 2,000 highly variable genes (HVGs) + kNN; (iii) HVG + PCA(50) + kNN; (iv) HVG + UMAP(50) + kNN (UMAP-learn v0.5, n_neighbors = 15, min_dist = 0.1) (30); (v) all genes + PCA(121) + kNN, matching the VAE latent dimensionality.

### Zero-shot benchmarks

scGPT (9) (human pre-trained checkpoint, 33M-cell pre-training) was applied in zero-shot mode: bulk RNA-seq samples were z-scored to scGPT’s input convention, embedded through the frozen model (768-dim CLS embedding), and classified by kNN (k = 5, cosine). No fine-tuning was performed; the result is therefore a measure of out-of-distribution transfer from single-cell to bulk RNA-seq.

BulkFormer is a recently published foundation model for bulk transcriptomes (12). It has ∼150 million parameters covering ∼20,000 protein-coding genes, pretrained on over 500,000 human bulk RNA-seq profiles. It combines a graph neural network, capturing gene–gene interactions, with a Performer attention module modelling global expression dependencies. We applied the published checkpoint in zero-shot mode: TARGET samples were embedded through the frozen model and classified by kNN (k = 5) against the embedded test-set reference, the same zero-shot protocol used for scGPT. Because this pretraining corpus is drawn from GEO and ARCHS4, it likely includes the TARGET samples used here for evaluation; the task is therefore not out-of-distribution for BulkFormer, as it is for our VAE.

### TARGET cross-dataset validation

TARGET paediatric cancer RNA-seq (n = 734, seven cancer types) was obtained from UCSC Xena (15,16). Expression values were aligned to the 16,115 model genes and harmonized by per-gene z-score normalization followed by rescaling to match the training per-gene mean and standard deviation. Samples were encoded by the trained Standard VAE and classified by kNN (k = 5, Euclidean) against the 118K reference embeddings. Expected tissue assignments were defined from established developmental biology: leukaemias → blood / bone marrow / lymphoid (haematopoietic origin); neuroblastoma → brain / nerve / adrenal gland (neural-crest origin); Wilms tumour and rhabdoid tumour of the kidney → kidney / ovary / uterus / testis (intermediate-mesoderm origin).

### Software and code availability

All scripts and figures generated for this work are deposited at GitHub (https://github.com/BIMSBbioinfo/flexynesis_tissue_vae_manuscript), built on the Flexynesis framework (21) using PyTorch (31) and PyTorch Lightning (32). The HDF5 dataloader, training scripts, and analysis pipelines are available in the same repository. Trained model weights, the curated 118K compendium in HDF5 format, and pre-computed embeddings are deposited at Zenodo (https://doi.org/10.5281/zenodo.20595537). A lightweight interactive demo is available at Hugging Face (https://huggingface.co/spaces/akalinLab/flexynesis-tissue-vae). The full model weights, training data, and code for exact inference are deposited at Zenodo and GitHub as described above.

## Notes

https://github.com/BIMSBbioinfo/flexynesis_tissue_vae_manuscript

### Competing Interest Statement

The authors have declared no competing interest.

