## Supplementary file for "An atlas-scale generative model for unified representation learning of bulk RNA-seq data"

**Table S1.** Comprehensive 75K-baseline vs 118K Scope B comparison. All metrics evaluated on each version's held-out test split (75K: n = 11,340; v3: n = 28,274). Bold indicates 118K Scope B improvements. pp = percentage points.

| **Metric** | **the 75K baseline (75K)** | **118K (this work)** | **Δ** |
| --- | --- | --- | --- |
| **Sample count (train)** | 63,777 | **118,263** | +85% |
| **Sample count (test)** | 11,340 | **28,274** | +149% |
| **UBERON tissue categories** | 43 | 42 | — |
| **Cell lines included** | Yes | **No (curated)** | — |
| **Class balancing** | None | **Stratified 10K cap** | — |
| **Training time (RTX 4060)** | ~3 h | ~5 h | — |
| **Peak RAM (data loading)** | 50+ GB CSV | **25 GB HDF5** | −50% |
| **Balanced accuracy (Standard)** | 90.7% | **94.9%** | +4.2 pp |
| **Weighted F1 (Standard)** | 93.7% | **96.2%** | +2.5 pp |
| **Per-gene ρ (Standard)** | 0.914 | **0.935** | +1.8% |
| **Per-gene ρ (Denoising)** | 0.919 | **0.934** | +1.5% |
| **Per-sample ρ (Standard)** | 0.766 | **0.829** | +6.3 pp |
| **Per-sample ρ (Denoising)** | 0.773 | **0.830** | +5.7 pp |
| **Imputation @ 10% (Denoising)** | 0.852 | **0.880** | +2.8 pp |
| **Imputation @ 20% (Denoising)** | 0.853 | **0.879** | +2.6 pp |
| **Imputation @ 30% (Denoising)** | 0.855 | **0.879** | +2.4 pp |
| **LISI / max** | 1.09 / 4.0 | 1.02 / 3.0 | purer |
| **kNN source mixing (k=20)** | 0.047 | **0.012** | −74% |
| **TARGET overall accuracy (n=734)** | 86.6% | 84.6% | −2.0 pp |
| **AML (n=196)** | 97% | **99%** | +2 pp |
| **ALL (n=194)** | 65% | **96%** | +31 pp |
| **AML Induction Failure (n=32)** | 94% | **100%** | +6 pp |
| **Neuroblastoma (n=162)** | 99% | 72% | −27 pp |
| **Wilms Tumor (n=132)** | 95% | 65% | −30 pp |
| **Rhabdoid Tumor (n=5)** | 80% | **100%** | +20 pp |
| **scGPT zero-shot (frozen)** | 61.0% | 61.0% | — |
| **All genes + kNN baseline** | 89.6% | **92.6%** | +3.0 pp |
| **Top-2000 HVG + kNN baseline** | 89.0% | **93.4%** | +4.3 pp |
| **All genes + PCA(121) + kNN baseline** | 87.4% | **92.0%** | +4.6 pp |
| **Supervised VAE advantage (vs all-kNN)** | +1.1 pp | **+2.3 pp** | wider |

**Table S2. Developmental-origin rubric used to score TARGET tissue-of-origin transfer. For each paediatric cancer type, the expected adult UBERON tissue categories were defined a priori from its established developmental lineage, with the supporting reference. A prediction was scored correct if the assigned tissue fell within the expected set. The mappings reflect canonical developmental biology rather than post-hoc selection; alternative groupings at the germ-layer level (e.g. collapsing all intermediate-mesoderm derivatives) yield the same qualitative conclusions.**

| **Cancer type** | **Developmental origin** | **Expected adult tissue categories** | **Reference** |
| --- | --- | --- | --- |
| Acute myeloid leukaemia | Haematopoietic / myeloid progenitors (bone marrow) | blood, bone marrow, lymphoid | Bonnet & Dick, Nat Med 1997 |
| Acute lymphoblastic leukaemia | Haematopoietic / lymphoid progenitors | blood, bone marrow, lymphoid | Bonnet & Dick, Nat Med 1997 |
| AML, induction-failure subtype | Haematopoietic / myeloid progenitors | blood, bone marrow, lymphoid | Bonnet & Dick, Nat Med 1997 |
| Neuroblastoma | Developing sympathetic nervous system (neural crest) | brain, nerve, adrenal gland | Maris, N Engl J Med 2010 |
| Wilms tumour | Intermediate mesoderm (precursors of kidney, uterus, ovary, testis) | kidney, ovary, uterus, testis | Hohenstein et al., Genes Dev 2015 |
| Kidney, rhabdoid tumour | Renal / metanephric mesenchyme | kidney | Hohenstein et al., Genes Dev 2015 |
| Clear cell sarcoma of the kidney | Renal mesenchyme (no well-defined adult correlate) | kidney | Hohenstein et al., Genes Dev 2015 |

*References: Bonnet D, Dick JE. Nat Med 1997;3:730-7. Maris JM. N Engl J Med 2010;362:2202-11. Hohenstein P, Pritchard-Jones K, Charlton J. Genes Dev 2015;29:467-82.*

**Table S3. Per-category tissue-of-origin accuracy on the TARGET cohort: supervised VAE versus BulkFormer.** Both models were evaluated out of the box on 734 TARGET paediatric tumours using the same k-nearest-neighbour protocol and developmental-expectation rubric (Table S2; Figure 6). Accuracy is the fraction of samples assigned to the expected adult tissue(s) of origin. BulkFormer was pretrained on GEO/ARCHS4 and may have encountered TARGET, so the transfer is out-of-distribution only for the supervised VAE.

| **TARGET cancer category** | **Expected tissue of origin (developmental)** | **Supervised VAE** | **BulkFormer** |
| --- | --- | --- | --- |
| Haematopoietic (AML, ALL) | Blood, bone marrow, lymphoid | >97% | >97% |
| Neuroblastoma | Brain, nerve, adrenal gland | 72.2% | 75.9% |
| Wilms tumour | Intermediate mesoderm (kidney, ovary, uterus) | 65.2% | 13.6% |
| Clear cell sarcoma of the kidney | Kidney (no clear adult correlate) | 0% | 0% |
| **Overall (734 samples)** | — | **84.6%** | **76.8%** |

### Supplementary Figures

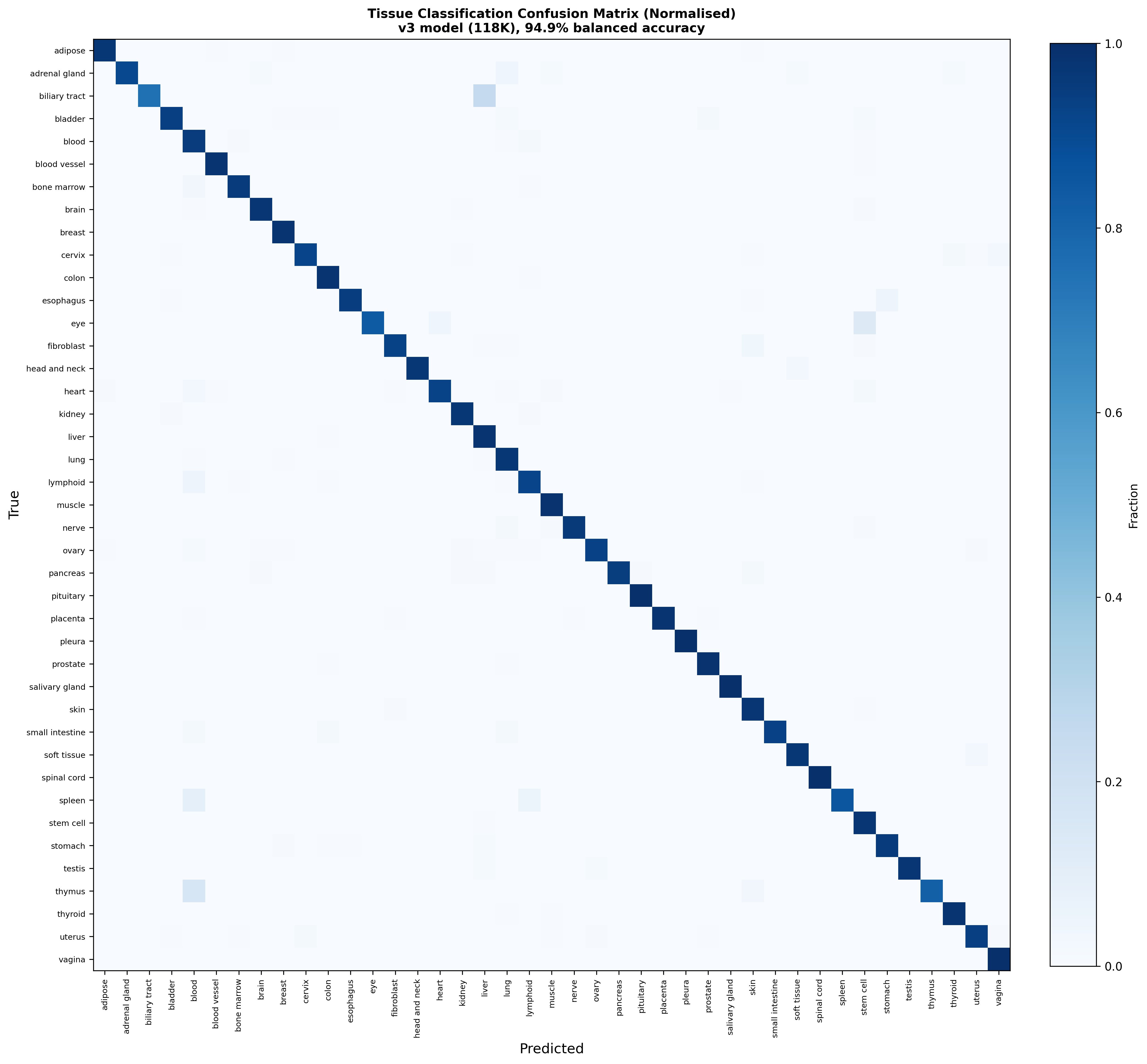

**Figure S1. Normalized confusion matrix for the 118K supervised VAE classifier.** Row-normalized confusion matrix across the 42 UBERON tissue categories on the held-out 118K test split (n = 28,274). Rows are true tissue labels, columns are predicted labels; diagonal values are per-class recall (matching the per-tissue accuracies in Figure 2). Off-diagonal entries highlight the few biologically plausible confusions remaining after class balancing.

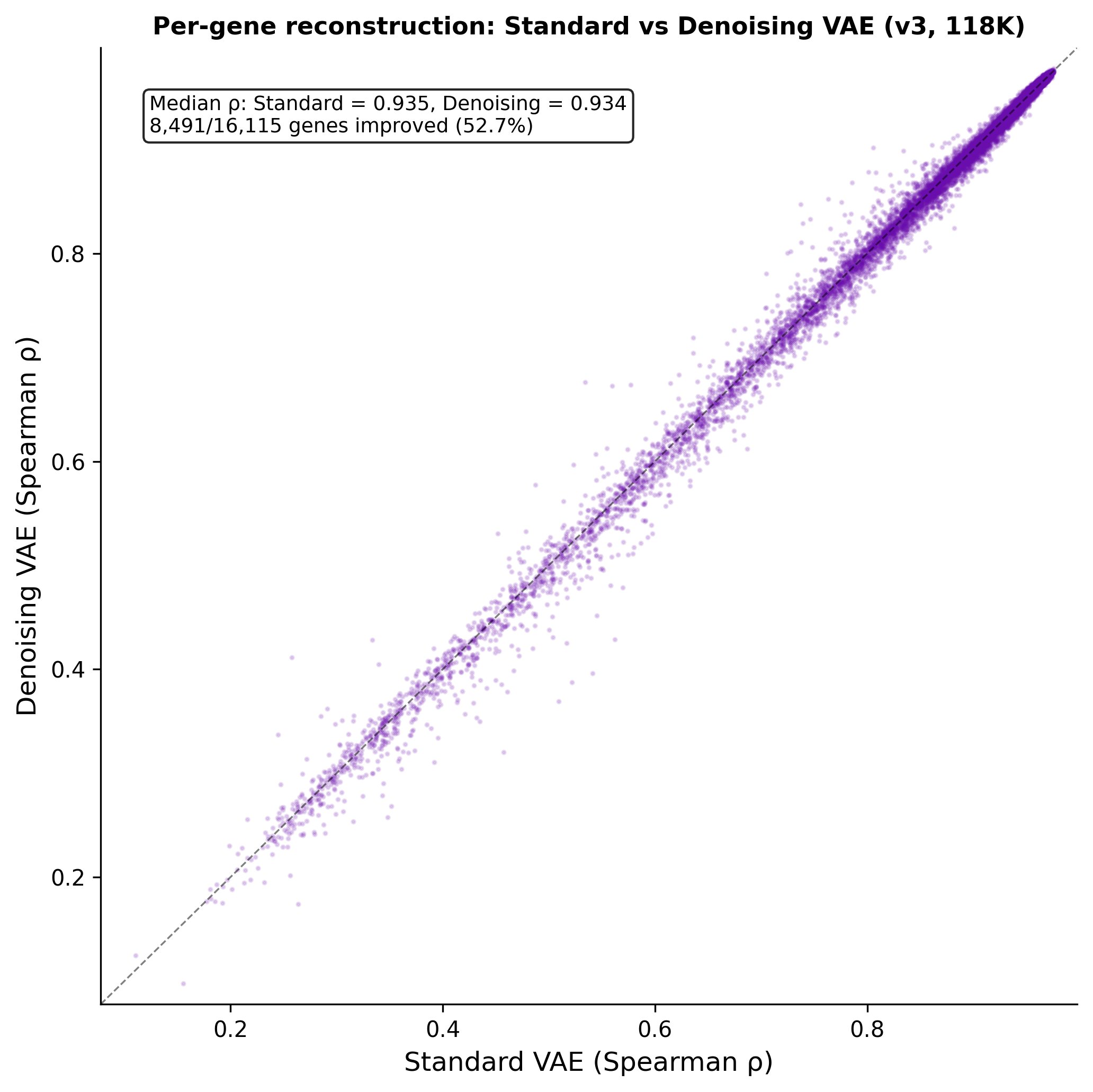

**Figure S2. Per-gene reconstruction performance scatter.** Scatter of original versus reconstructed expression values, aggregated across all genes and samples in the held-out 118K test split, for the Standard VAE variant. Density coloring shows the high-density diagonal consistent with the median per-gene Spearman ρ = 0.935 reported in Figure 3a.

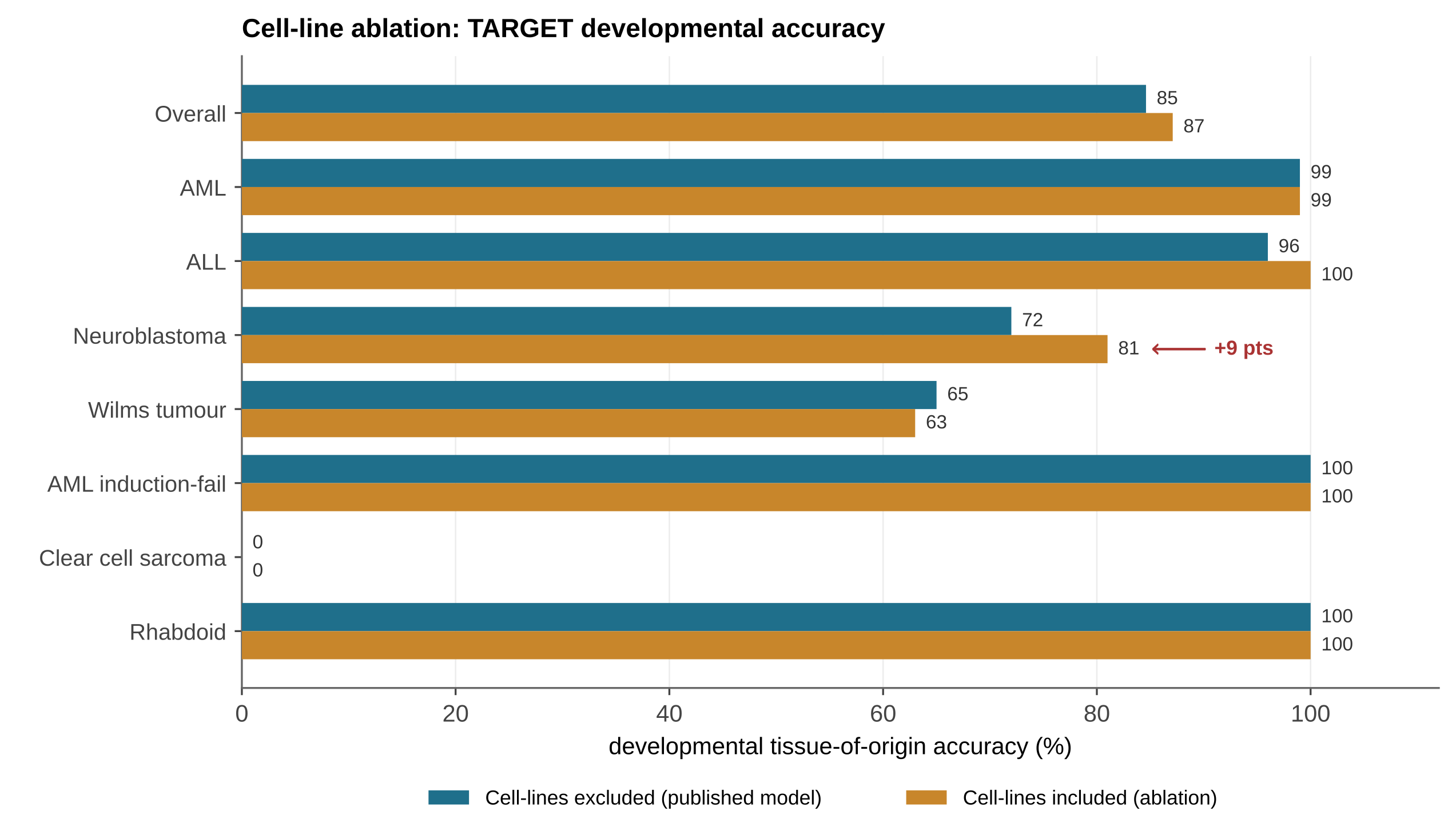
**Figure S3. Cell-line removal has a small, localized effect on TARGET developmental transfer.** Per-cancer and overall accuracy for the published model (cell lines excluded, teal) versus a matched control in which ARCHS4 cell lines were retained under the same per-tissue capping policy and random seed (cell lines included, ochre); the two models differ only in cell-line inclusion. Excluding cell lines changes overall accuracy by 2.5 percentage points (87.1% with cell lines, 84.6% without), an effect confined almost entirely to neuroblastoma (72% versus 81%), the one cancer whose neural-crest lineage cell lines most directly represent. All other cancer types are unchanged, and Wilms tumour differs by under two points (65% versus 63%). Cell-line removal therefore carries only a minor, lineage-specific cost on developmental transfer, consistent with its role in producing a latent space less anchored to cell-line transcriptomes (Table 2).
